# geoGraph: Connectivity modelling across geographic space with hexagonal discrete global grids

**DOI:** 10.64898/2026.10.14.759071

**Authors:** Dominik C. Jud, Evelyn J. Carter, Andrea Vittorio Pozzi, Ondrej Pelanek, Aramish Fatima, Margherita Colucci, Cecilia Padilla-Iglesias, Michela Leonardi, Shujie Chang, Cassandra Gunasekaram, Lucio Vinicius, Andrea Bamberg Migliano, Thibaut Jombart, Andrea Manica

**Affiliations:** Human Evolutionary Ecology Group, Department of Evolutionary Anthropology, University of Zürich, Zürich, Switzerland; Evolutionary Ecology Group, Department of Zoology, University of Cambridge, Downing Street, Cambridge, CB2 3EJ, UK; Faculty of Science and Engineering, Life Sciences, Anglia Ruskin University, Cambridge, UK; University Museum of Zoology, University of Cambridge, Downing Street, Cambridge, CB2 3EJ, UK; Human Palaeosystems Group, Max Planck Institute of Geoanthropology, Jena 07743, Germany; HUMAN ORIGINS – Cluster of Excellence for Integrative Human Origins Studies (EXC 3101), University of Tübingen, Tübingen, Germany; Natural History Museum, London, SW7 5BD, UK; Deep-time Ecology Group, Department of Zoology, University of Cambridge, Downing Street, Cambridge, CB2 3EJ, UK; MRC Centre for Global Infectious Disease Analysis, School of Public Health, Imperial College London, UK

**Keywords:** geographic information systems, landscape connectivity, landscape genetics, least-cost path, population genetics, R package, spatial graphs

## Abstract

1. Estimating the movement and connectivity of organisms across landscapes is central to research in ecology and evolutionary biology. Although graph-based approaches representing geographic space as weighted networks provide a flexible and computationally efficient framework for such analyses, existing tools either perform their computations only on rectangular raster grids, thus introducing systematic directional and area distortions at global scales, or rely on bespoke pipelines that limit the general applicability of recent hexagonal-grid analyses.
2. To overcome these shortcomings, here we present *geoGraph*, an R package that brings graph-based spatial modelling to discrete global hexagonal grids. By representing landscapes on an equal-area hexagonal tessellation of the sphere, *geoGraph* enables global-scale analysis without projection distortion while retaining the flexibility of the graph framework and the performance of the graph and RBGL/Boost libraries. The package integrates both raster- and vector-based GIS data into a single cost surface, supports custom grids at various resolutions, and provides a complete pipeline from raw GIS layers to graph-algorithmic outputs, together with tools for visualising and interactively editing graphs.
3. We illustrate *geoGraph* by revisiting previous studies of isolation by distance among human populations in the Human Genome Diversity Project panel. At a global scale, restricting movement to land raised the correlation between geographic and genetic distance from r = 0.74 (great-circle) to r = 0.84. On a finer grid within Asia, accounting for terrain ruggedness improved the fit, with the best-fitting models implying reduced movement across the Himalaya and Tibetan Plateau. These analyses demonstrate that *geoGraph* can model movement and connectivity appropriate to the desired spatial scale and that the level of geographic realism required is itself scale-dependent.
4. By combining discrete global grids with the power of graph algorithms, *geoGraph* enables seamless connectivity analysis from landscape to global scales. It is broadly applicable across landscape genetics, phylogeography, conservation planning, and studies of human dispersal, and is particularly suited to large-scale problems that rectangular-raster tools cannot address without distortion.

## 1 Introduction

Estimating the movement and connectivity of organisms is an essential part of many studies in ecology and evolutionary biology (Manel et al., 2003; Taylor et al., 1993). Ever since high-resolution spatial data became widely available, researchers have increasingly relied on spatially explicit models to study dispersal and connectivity across landscapes (Calabrese & Fagan, 2004; Kozak et al., 2008; Manel & Holderegger, 2013). Unlike straight-line Euclidean distance, movement across a heterogeneous landscape better represents geographic separation and population isolation (Landguth et al., 2010; McRae, 2006; Wang et al., 2013). Graph-based approaches have long provided a natural and mathematically rigorous framework for such analyses, representing the landscape as a network of nodes connected by weighted edges (Urban & Keitt, 2001). Because landscapes can be discretised at any chosen resolution and edge weights can encode arbitrary cost rules, this framework accommodates everything from individual habitat patches to continent-scale grids and allows for efficient computation of shortest paths, connectivity metrics, and patterns of accessibility across heterogeneous terrain. This flexibility is why graphs have become widely used across landscape ecology and conservation planning (Pascual-Hortal & Saura, 2006; Rayfield et al., 2011), population and landscape genetics (Dyer & Nason, 2004), and phylogeography and human dispersal (Manica et al., 2005; Prugnolle et al., 2005).

A crucial preliminary step in graph-based approaches involves transforming geographic data into a graph representation that preserves spatial structure, encodes relevant environmental attributes, and allows movement costs to be defined in a flexible and transparent manner. However, interfacing spatial data with graph-based methods can be a complicated task. Spatial datasets are typically stored as raster objects, whereas graph algorithms operate on abstract graph structures, so bridging these two representations often requires substantial custom code and careful bookkeeping.

Several tools address parts of this workflow (Dutta et al., 2022). The *gdistance* package (Van Etten, 2017) computes cost-distance and randomised shortest-path distances on grids. Based on this, the *graph4lg* package (Savary et al., 2021) builds and compares landscape and genetic graphs for landscape genetics applications. The *topoDistance* package (Wang, 2020) computes topographic distances between raster cells. *Circuitscape* (McRae et al., 2016) models connectivity using circuit theory. Importantly, most of these tools operate on rectangular raster grids. While some apply geometric corrections for grid distortion, such as *gdistance* (Van Etten, 2017), no rectangular grid can be made equal-area and free of directional bias across the globe, so residual area and directional distortion arise when used at global scales (Battersby et al., 2017; Gudmundsson & Alerstam, 1998). Discrete global grids avoid these distortions by tessellating the sphere directly into cells of approximately equal area and shape (Sahr et al., 2003). While recent work has begun to move to such grids for large-scale analyses, these largely rely on bespoke pipelines rather than reusable infrastructure. Tools such as GAIA (Grundler et al., 2025) and CISGeM (Eriksson et al., 2012; Padilla-Iglesias et al., 2026), for example, can achieve realistic migration costs using movement friction surfaces based on discrete global grids (Lian & Python, 2026), yet the grids and friction surfaces they require are typically built with custom code. In parallel, methods such as EEMS (Petkova et al., 2016), MAPS (Al-Asadi et al., 2019) and FEEMS (Marcus et al., 2021) estimate migration surfaces from genetic data across grids of demes, but these estimates are usually interpreted on their own rather than reused in further spatial models.

Here, we present *geoGraph* v2.0.1, an R package (R Core Team, 2021) that brings discrete global hexagonal grids to graph-based spatial modelling. By representing landscapes on a hexagonal tessellation of the sphere rather than on rectangular rasters, *geoGraph* allows for global-scale analysis without projection distortions while retaining the algorithmic power and flexibility of the graph framework. *geoGraph* models movement and connectivity across spatially explicit landscapes and enables the integration of raster-based data and spatial polygon (vector) data into a single cost surface, providing a complete pipeline from raw GIS layers to graph-algorithmic output. Because grids and cost surfaces are stored as standard graph objects, they can also serve as inputs to friction-based dispersal models. Conversely, migration surfaces estimated from genetic data share this graph structure of connected demes and could likewise be imported into *geoGraph* as movement costs. Together, this makes graph-based connectivity analysis accessible and easy to use even at large spatial scales.

## 2 Design and implementation

The *geoGraph* package provides classes and methods for representing geographic space as a graph and analysing movement, dispersal, and connectivity across it. Geographic regions are modelled by a regular grid, where each node has a set of spatial coordinates and a set of attributes, such as habitat descriptors or the presence/abundance of a given species. Travelling within a region is then modelled simply as moving between connected nodes, and the cost of moving from one node to another can be defined from their attribute values. This allows movement costs to range from binary land/sea connectivity to multi-layer environmental cost surfaces derived from raster and vector GIS data. Once a graph has been built, *geoGraph* harnesses the full power of graph algorithms implemented in R by the *graph* (Gentleman et al., 2026) and *RBGL* (R Boost Graph Library) (Carey et al., 2026) packages. In particular, *RBGL* is an interface between R and the comprehensive Boost Graph Library in C++ (Siek et al., 2001), which provides fast and efficient implementations of a wide range of graph algorithms. *geoGraph* also provides functions for visualising graphs, paths, and node attributes on geographic maps, and for interactively editing the graph structure to correct artefacts or incorporate biological knowledge.

### 2.1 Data representation

Data in *geoGraph* is stored in two distinct types of objects, *gGraph* and *gData*, providing a clear separation between the representation of a spatial landscape and the representation of data or locations embedded within that landscape (Fig. 1). Both object types are defined as formal (S4) classes and often have distinct methods for the same generic function (e.g. *getNodes()* is defined for both classes). *gGraph* objects represent the spatial structure itself, including a spatial grid and node attributes. On the other hand, *gData* objects represent sets of locations (e.g. sampling sites or observation points) that have been interfaced with a *gGraph*, enabling subsequent graph-based operations such as computing paths or distances between locations. This functional separation ensures that landscape structure and empirical data can be modified independently, improving reproducibility and flexibility (Alston & Rick, 2021).

**Figure 1:**
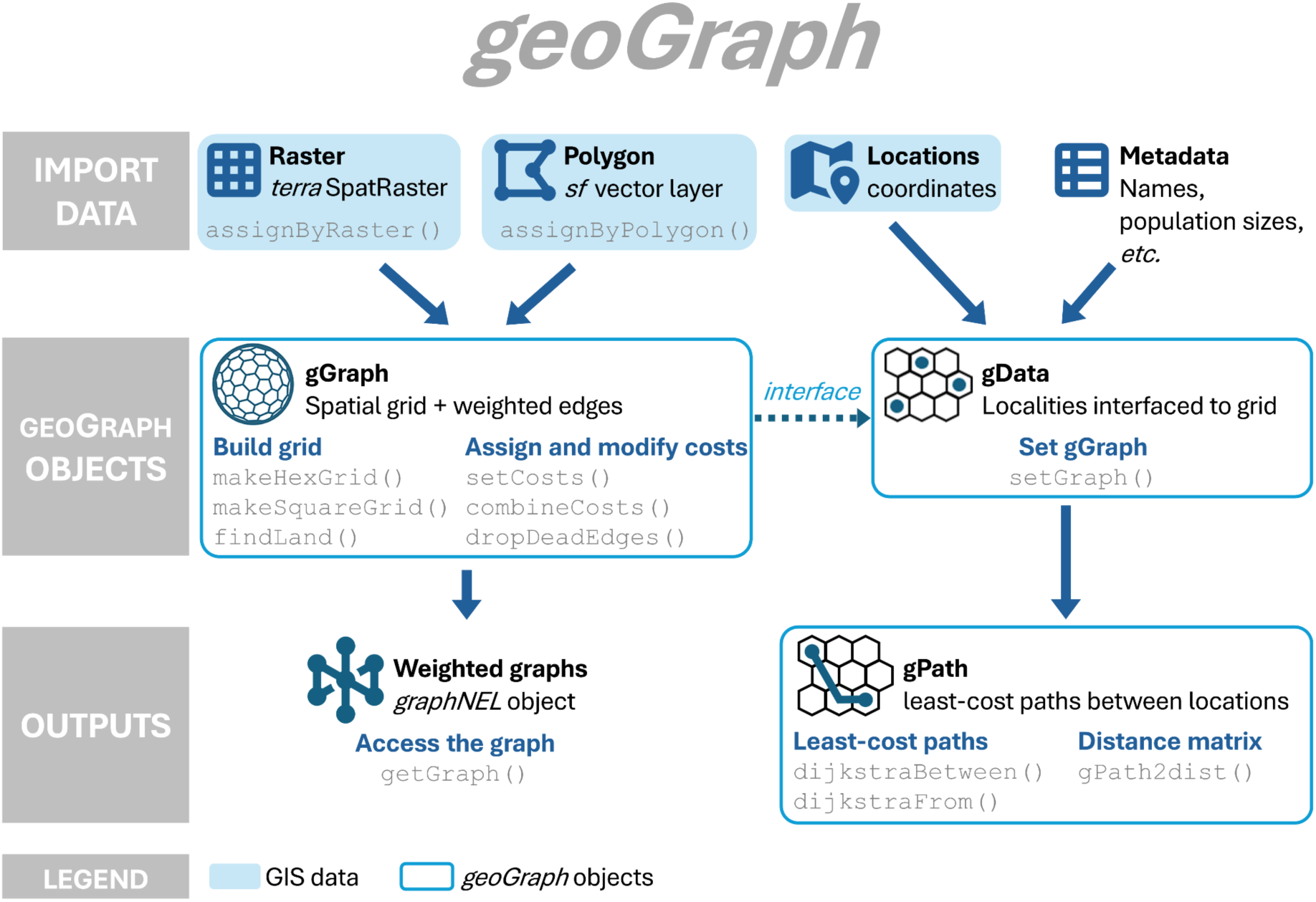
Schematic *geoGraph* workflow. Standard spatial data are imported into *geoGraph’s* two formal (S4) classes. For *gGraphs*, node attributes can be assigned either by raster layers (*terra* SpatRaster) or vector polygons (*sf*) with the functions *assignByRaster()* and *assignByPolygon()* respectively. Based on these node attributes, edge weights (costs) can be set. Sample locations and their associated metadata are then interfaced with a *gGraph* into a *gData* on which *geoGraph* computes least-cost paths and pairwise distances.

#### 2.1.1 gGraph objects

The graph component underlying the *gGraph* is a *graphNEL* object, the standard graph class in the *graph* and *RBGL* packages. It stores all information on the connections between nodes (formally the graph edges), and their weights (costs). In addition to the graph structure, *gGraph* objects store spatial coordinates of the nodes and their auxiliary information (e.g. habitat, temperature).

Several examples of *gGraph* objects are distributed with *geoGraph,* namely the *rawgraph.10k, rawgraph.40k, worldgraph.10k,* and *worldgraph.40k*. These grids are custom-built based on the Icosahedral Snyder Equal Area (ISEA) projection (Snyder, 1992), which partitions the globe into cells of approximately equal area. As this avoids the distortion of longitude-latitude grids toward the poles, ISEA-based discrete global grids are among the best suited to global-scale statistics (Kmoch et al., 2022). The difference between *rawgraphs* and *worldgraphs* is that in the former all adjacent cells are connected, while in the latter connections occur only on land. The numbers 10k and 40k indicate that the grids consist of roughly 10,000 and 40,000 nodes, respectively. These included *gGraph* objects carry deliberately simple habitat information on a relatively coarse grid, but richer descriptors can be added by extracting more attributes from GIS layers (Section 2.3), and a finer grid can be built as a custom *gGraph* (Section 2.2).

#### 2.1.2 gData objects

*gData* objects represent sets of locations that have been interfaced with a *gGraph* object. They provide the link between empirical spatial data (e.g. sampling sites, populations, observation points) and the graph-based representation of geographic space implemented in *geoGraph*. Constructing a *gData* object involves assigning each location to the closest node on the underlying *gGraph* grid. This interfacing step establishes a correspondence between original geographic coordinates and graph nodes, which then serve as the entry points for all subsequent graph-based analyses. Once locations have been assigned to a node, movement and connectivity can be modelled, allowing distances and least-cost paths to be computed, all under alternative movement-cost scenarios encoded in the *gGraph*. *gData* objects can also store additional information, such as population sizes or species presence data, and provide access to the attributes of the underlying *gGraph* at the matched nodes. This allows environmental or habitat information stored on the grid to be directly associated with locations, and subsequently used to define movement costs or to compare alternative landscape models.

### 2.2 Creating a custom graph

While the prebuilt global grids cover most large-scale applications, *geoGraph* also provides functions to construct custom spatial graphs. The function *makeHexGrid()* creates a discrete global hexagonal grid using the R package *dggridR* (Barnes & Sahr, 2026), which builds on DGGRID (Sahr, 2024). The grid is based on the ISEA projection (Snyder, 1992), so cell areas are approximately equal everywhere and inter-node distances do not vary with latitude, two properties that are important for unbiased least-cost path computation.

The user can specify a geographical bounding box and an inter-node spacing, which can range from over 4,000 km down to just above 100 m. From these, *geoGraph* takes each cell centre of the hexagonal grid as a node and each connection between neighbouring centres as an edge, producing a raw *gGraph* and its underlying *graphNEL*. Because the number of cells grows with the area covered and inversely with the inter-node spacing, fine grids over large extents can yield very large graphs that are memory- and time-intensive to build and analyse. The spacing should therefore be chosen to balance the resolution required against computational cost.

For smaller extents, the *makeSquareGrid()* function creates a traditional rectangular grid with rook-style (four-neighbour) connections over a specified bounding box at a chosen resolution. This is suited for quick analyses over small geographic areas where projection distortion is negligible.

Custom *gGraph* objects have uniform movement costs after grid creation, but their edges can be weighted from node attributes to reflect the landscape (Section 2.3).

### 2.3 Assigning environmental attributes and defining costs

A defining feature of *geoGraph* is the ability to populate grid nodes with environmental attributes and use them to define movement costs. This is handled by the *nodes.attr* slot of a *gGraph* where each node can be assigned several attribute values. For example, after creating a new raw *gGraph*, the function *findLand()* assigns land or sea status to each node by spatial overlay with a global coastline layer from *rnaturalearth* (Massicotte & South, 2026), populating the *nodes.attr* slot with a ‘habitat’ column. Another way to add node attributes is by adding data from any standard GIS data type using the functions *assignByRaster()* for raster data and *assignByPolygon()* for vector data.

For example, if the user wants to add data from an elevation raster, the function *assignByRaster()* aggregates the cells from the raster to the nodes in a *gGraph* by assigning each cell to the closest node. As this results in multiple cells being assigned to the same node, the user can summarise the raster values per node with any user-specified function. In the case of the elevation raster, the standard deviation of elevation values assigned to each node, for instance, provides a per-node measure of topographic ruggedness.

Alternatively, the function *assignByPolygon()* extracts attributes from any *sf* polygon layer, such as a biome classification, and assigns them to nodes by spatial intersection. For example, supplying a polygon of the Sahara desert labels every node that falls within it with the attribute ‘desert’. Multiple attributes from different sources can be combined in the *nodes.attr* data frame, and new derived attributes can be added at any time using *setNodesAttr()*.

Once node attributes have been assigned, edge weights are set using *setCosts()*, which computes the weight of each edge from the attribute values of its two endpoint nodes. The *method* argument controls how the two endpoint values are combined (by default their mean, or any user-supplied binary function). Specific cost values for each attribute level are defined in a cost rules table stored in the *gGraph’s* meta slot, and can be updated at any time. Cost surfaces from multiple environmental layers can be composed using *combineCosts()*, which multiplies or adds per-node cost contributions from different attributes. Edges whose cost exceeds a threshold can be removed with *dropDeadEdges()*, effectively restricting movement to permissible habitat types (e.g. land-only or sea-only connectivity).

### 2.4 Editing and visualising graphs

*geoGraph* also lets users refine a *gGraph* object by hand, either by locally adding or removing connections, or by altering the attributes of individual nodes. This may be needed to connect components such as islands to the mainland, or to correct artefacts introduced during grid construction or data extraction. Such local edits provide fine-grained control over the graph structure and enable the incorporation of more detailed knowledge.

When customising a *gGraph* object, it is often useful to visually inspect specific regions, and more generally to navigate the graphical representation of the data. This can be done by using the interactive functions *geo.zoomin()*, *geo.zoomout()* and *geo.slide()* directly called by clicking on a plotted *gGraph* object. In order to provide these interactions, the standard *plot()* and *points()* functions use base R graphics, as they rely on the *locator()* function to capture mouse clicks on the active plotting device.

Plotting a *gGraph* displays the grid of nodes over a background shapefile (by default, landmasses), and edges can be shown at the same time via the *edges* argument, or added afterwards with *plotEdges()*. Because a *gGraph* may contain a very large number of nodes and edges, plotting can be demanding on some graphical devices, hence it is advisable to restrict visualisation to specific regions or subsets when exploring large grids. *gData* objects are by default plotted on top of their associated *gGraph*, so that locations can be seen in their spatial and graph contexts and compared directly with their corresponding grid positions.

For more customisable, publication-ready visualisation, *geoGraph* also provides *autoplot()* methods that render *gGraph* and *gData* objects using *ggplot2* (Wickham, 2016). Because these return standard *ggplot* objects, they can be extended with any *ggplot2* layer, scale or theme, making it easy to produce polished figures or drop *geoGraph* output into existing *ggplot*-based workflows.

## 3 Example Applications

One of the most useful applications of *geoGraph* is the computation of least-cost paths between pairs of locations based on Dijkstra’s (1959) algorithm. Least-cost paths are a widely used approach for modelling dispersal routes and effective distances across heterogeneous landscapes (Van Etten & Hijmans, 2010; Wang et al., 2009). This can be achieved using the functions *dijkstraFrom()* and *dijkstraBetween()* on a *gData* object which contains all the locations of interest. These functions return the paths as *gPath* objects. *dijkstraFrom()* computes the paths from a given node of the grid to all locations of the *gData*, while *dijkstraBetween()* computes the paths between pairs of locations of the *gData*.

### 3.1 Worldwide least-cost paths

To illustrate these functions and their integration with genetic data, we revisit a classical result in human population genetics, the isolation-by-distance signal in the Human Genome Diversity Project (HGDP) panel (Cann et al., 2002), where pairwise genetic differentiation between human populations was shown to increase with geographic distance (Manica et al., 2005). We compared pairwise F_ST_ among 51 HGDP populations to two inter-population distances: great-circle distances between sampling coordinates, and least-cost path distances on the *worldgraph.40k*, where movement is restricted to land nodes (Fig. 2a). Both distance matrices were correlated with the F_ST_ matrix using a Mantel test (Mantel, 1967) with 999 permutations, as implemented in the *vegan* package (Oksanen et al., 2026). As expected, restricting movement to land markedly improved the fit over straight-line distance, with the Mantel correlation rising from r = 0.74 (great-circle distances, Fig. 2b) to r = 0.84 (land-bound least-cost paths, Fig. 2c). This simple comparison demonstrates how the deployment of geographically explicit distances via *geoGraph* clearly outperforms approaches based on straight-line distances at global scale (Handley et al., 2007). The 40,000-node *worldgraph* in *geoGraph* allows for flexible global-scale analyses without manually placed waypoints (Ramachandran et al., 2005).

**Figure 2:**
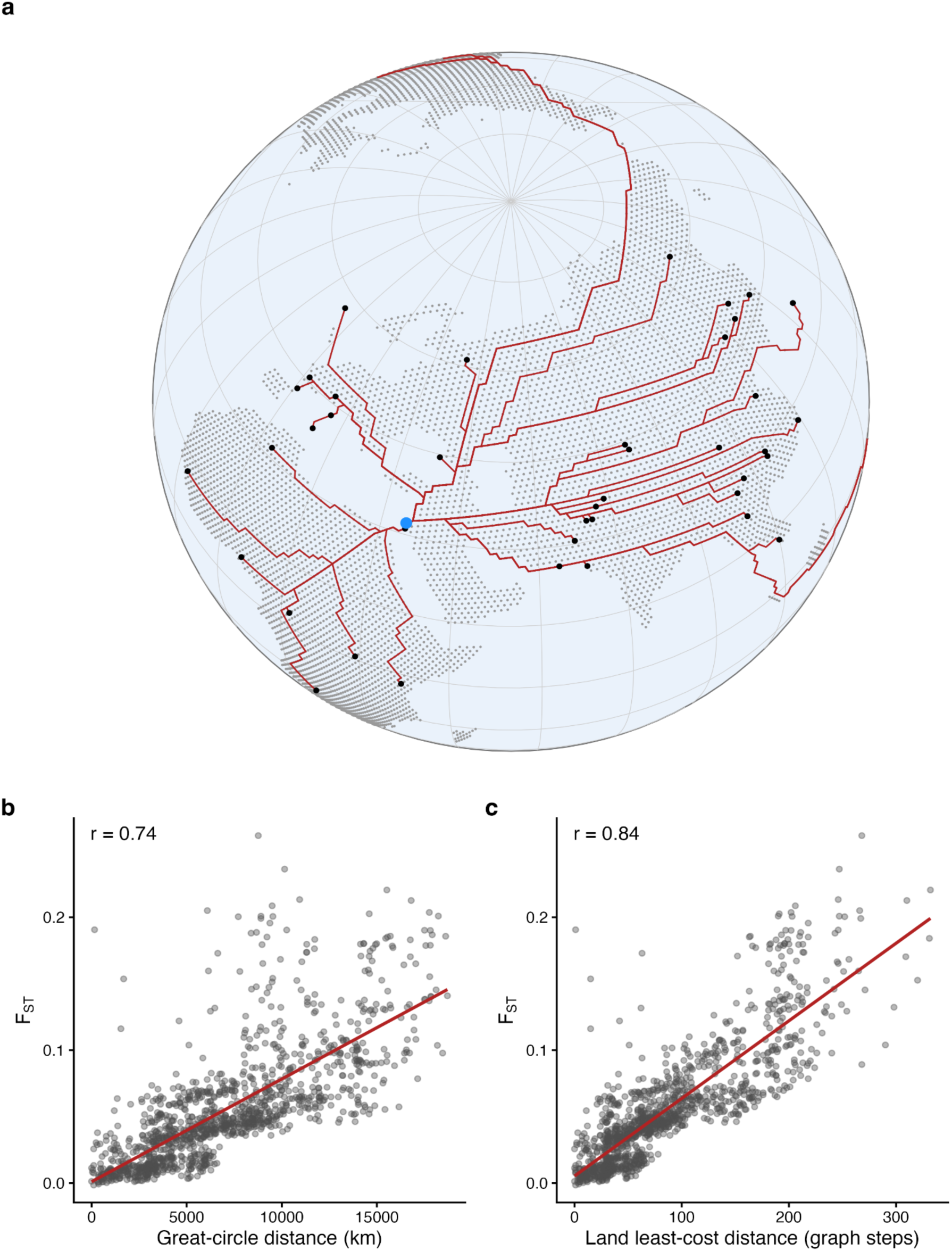
Global isolation by distance in the HGDP panel. (a) Land-restricted least-cost paths across the worldgraph.40k grid from one origin (Druze, blue) to all populations (black), computed with *dijkstraFrom()*. (b, c) Pairwise F_ST_ against great-circle distance (b) and land-bound least-cost distance in graph steps (c); points are population pairs, red lines are linear fits, r is the Mantel correlation. Constraining movement to land improves the fit from r = 0.74 to r = 0.84.

### 3.2 Building a custom hexagonal grid

While limiting movement costs to land may suffice at global scale, analyses within a single landmass can benefit from accounting for terrain ruggedness, which is known to shape movement and gene flow across topographically complex landscapes (Murphy et al., 2010; Wang et al., 2013). We therefore restricted our second analysis of correlations between genetic and geographical distances to the 25 continental Asian HGDP populations and built a custom hexagonal grid over Asia with *makeHexGrid*() at ∼50 km inter-node spacing (Fig. 3a). Using *assignByRaster()*, we assigned each node the standard deviation of elevation within its cell as a simple measure of topographic ruggedness (cf. Riley et al., 1999). Elevation data were obtained from the *elevatr* package (Hollister et al., 2023) at zoom level 4 and clipped to the study region, yielding a raster of 0.037° resolution (≈4 km). Edge costs were then derived from these values with *setCosts()*, and edges whose ruggedness exceeded a threshold were removed with *dropDeadEdges()*, so that least-cost paths avoid the most rugged terrain, such as the Himalaya and Tibetan Plateau. Next, we estimated correlations between the obtained plain least-cost distances and F_ST_ for the 25 Asian populations.

**Figure 3:**
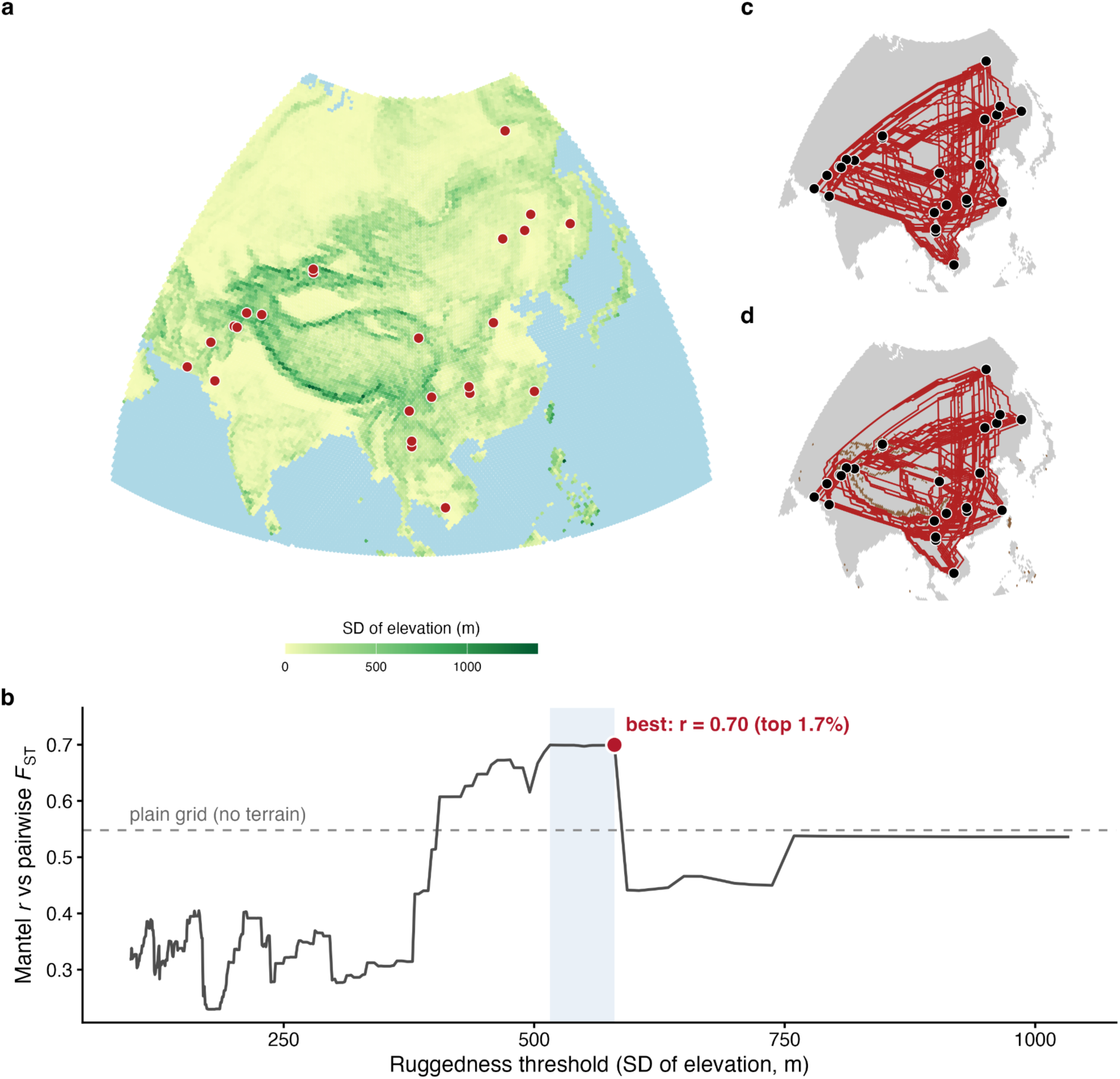
Terrain ruggedness and genetic differentiation in Asia. (a) Custom ∼50 km hexagonal grid over Asia coloured by ruggedness (SD of elevation within each cell), with the 25 HGDP populations (red); the Himalaya forms a high-ruggedness spine. (b) Exploratory sweep of the ruggedness threshold: Mantel r between pairwise F_ST_ and ruggedness-weighted least-cost distance, versus the no-terrain baseline (dashed, r ≈ 0.54); best fit r = 0.70 (plateau shaded). (c, d) Least-cost paths between populations without (c) and with (d) terrain cost, with a visible reroute around the mountains.

On this finer grid, correlations based on plain least-cost distances tracked great-circle distances closely (r = 0.54 and r = 0.55, respectively), reflecting that all populations lie on one connected landmass. To examine the influence of terrain, we used *geoGraph* to explore how the fit changed across ruggedness thresholds. This sweep is not a validated parameter estimate, but it illustrates how readily the package allows users to search the cost-parameter space. Fit improved over a broad range of thresholds that penalise only the most extreme terrain, namely the top ∼2-3% of ruggedness values, reaching r ≈ 0.70 (Fig. 3b). This regime isolates the Himalaya and Tibetan Plateau (Fig. 3c,d), consistent with these ranges acting as major barriers to gene flow between the South/Central and East Asian populations (Gayden et al., 2007; Zhang et al., 2022). As the fit was nonetheless sensitive to the exact threshold, we treat this result as exploratory rather than as a formal test.

## 4 Conclusions

*geoGraph* provides a complete and easily accessible tool for graph-based spatial modelling on equal-area discrete global grids. The package integrates both raster-and vector-based GIS data onto equal-area hexagonal grids, and allows the user to define and combine cost surfaces while retaining the full flexibility of the graph framework. *geoGraph’s* two-object design separates landscape structure (*gGraph*) from empirical data (*gData*), allowing for flexible and reproducible analysis within a single environment. By lowering the technical barrier to this kind of analysis, *geoGraph* offers reusable, general-purpose infrastructure for building and analysing spatial graphs from local to global scales.

Our examples illustrate that the appropriate level of geographic realism is scale-dependent. At global scale, restricting movement to land substantially improved the prediction of genetic differentiation among HGDP populations, whereas within Asia, where all populations share a single landmass, adding terrain ruggedness and switching to a finer grid improved the fit. These results highlight two capabilities that make *geoGraph* well suited to such analyses. First, because grid resolution matters, the ability to build custom grids at any resolution with *makeHexGrid()* allows users to match the grid to the process and scale of interest. Second, because movement costs can be parameterised in many ways and model fit can be sensitive to these choices, *geoGraph’s* integration of raster and vector layers into combinable cost surfaces makes it straightforward to define, compare, and explore alternative cost models.

Lastly, because computing time scales with the number of nodes and edges, very fine grids over large extents can be memory-and time-intensive. In practice, however, grids of a few tens of thousands of nodes are sufficient for many continental-to global-scale questions and run quickly on a standard desktop. Because *geoGraph* reads standard *terra* and *sf* objects, it interoperates directly with the wider R spatial ecosystem, enabling cost surfaces to be assembled from essentially any GIS data source. This also opens a route to temporally dynamic analyses: coupling *geoGraph* with palaeoclimate reconstructions such as those from *pastclim* (Leonardi et al., 2023) would allow movement costs to be recomputed through time, a promising direction for modelling past dispersal and range shifts.

More broadly, *geoGraph* is applicable wherever movement, dispersal, or connectivity must be modelled across geographic space, from landscape genetics and phylogeography to conservation planning and studies of human dispersal. By bringing the power and flexibility of graph algorithms to equal-area discrete global grids, it makes connectivity analysis straightforward from local to global scales.

## Acknowledgements

We thank François Balloux for feedback on the initial version of the package, and Anna Graff for her input and for testing package functions.

D.C.J. is funded by the NCCR Evolving Language (Swiss National Science Foundation agreement no. 51NF40_180888) and the A. H. Schultz Foundation. E.J.C. is supported by a Cambridge Philosophical Society Sedgwick studentship. A.V.P. and O.P. are supported by the Natural Environment Research Council (grant no. NE/S007164/1). A.F. is supported by the John Stanley Gardiner Studentship from the Department of Zoology, University of Cambridge. M.C. is supported by the Max Planck Society (Human Palaeosystems Research Group). C.P-I. is funded by an Alan Wilson Research Fellowship from Emmanuel College, Cambridge. M.L. was supported by the Leverhulme Research Grant RPG-2020-317 and the ERC Synergy Grant 101224035 “FORAGER”. C.G. is funded by the Swiss National Science Foundation (grant no. 10.001.620) and the A. H. Schultz Foundation. A.M. is supported by UK Research and Innovation (grant no. UKRI6329).

LLM use: During code development, Copilot was used for autocomplete-style suggestions in RStudio. On GitHub, Copilot was used for automatic code review of pull requests made by contributors, with human reviewers deciding which suggestions to implement.

## Author contributions

A.M. and T.J. designed the package functionality, with input from D.C.J. A.M., T.J., D.C.J., C.P-I., E.J.C., A.F., A.V.P., O.P., S.C., C.G., M.L. and M.C. wrote the R package and documentation. A.M., T.J., D.C.J., E.J.C., A.V.P., M.C. and C.P-I. wrote unit tests. D.C.J. led the writing of the manuscript, with input from A.M., M.L., A.V.P., S.C., C.P-I., L.V. and A.B.M. All authors contributed critically to the drafts and gave final approval for publication.

## Conflict of interest

The authors declare no conflicts of interest.

## Software availability

The package and its documentation are distributed under the free, open-source MIT licence, and can be downloaded from the following GitHub repository: https://github.com/EvolEcolGroup/geograph.

## Data availability

The HGDP data and global grids used in this study are distributed with *geoGraph*; elevation data were obtained via the *elevatr* package. Scripts to reproduce all analyses are available at https://github.com/EvolEcolGroup/jud_geograph_manuscript.

## Statement of Inclusion

Our study is methodological, revisiting a previously published, publicly available dataset (the Human Genome Diversity Project panel) to demonstrate a new R package; it involved no local or primary data collection. Because the contribution is a general-purpose tool rather than a study of any specific population, no region-specific fieldwork or local collaboration was required. We cite the original data generators and make our software and code openly available for reuse in any region.

## Notes

### Competing Interest Statement

The authors have declared no competing interest.

